# Effects of the glucocorticoid drug prednisone on urinary proteome and candidate biomarkers

**DOI:** 10.1101/128603

**Authors:** Jianqiang Wu, Xundou Li, Manxia An, Youhe Gao

## Abstract

Urine is a good source of biomarkers for clinical proteomics studies. However, one challenge in the use of urine biomarkers is that outside factors can affect the urine proteome. Prednisone is a commonly prescribed glucocorticoid used to treat various diseases in the clinic. To evaluate the possible impact of glucocorticoid drugs on the urine proteome, specifically disease biomarkers, this study investigated the effects of prednisone on the rat urine proteome. Urine samples were collected from control rats and prednisone-treated rats after drug administration. The urinary proteome was analyzed using liquid chromatography–tandem mass spectrometry (LC-MS/MS), and proteins were identified using label-free proteome quantification. Differentially expressed proteins and their human orthologs were analyzed with bioinformatics methods. A total of 523 urinary proteins were identified in rat urine. Using label-free quantification, 27 urinary proteins showed expression changes after prednisone treatment. A total of 16 proteins and/or their human orthologs have been previously annotated as disease biomarkers. After functional analysis, we found that the pharmacological effects of prednisone were reflected in the urine proteome. Thus, urinary proteomics has the potential to be a powerful drug efficacy monitoring tool in the clinic. Meanwhile, alteration of the urine proteome due to prednisone treatment should be considered in future disease biomarker studies.

## Introduction

Disease biomarkers are measurable changes associated with specific pathophysiological conditions that can be used to diagnose and monitor diseases, to predict prognoses and to reflect treatment efficiency. In contrast to blood, urine accumulates systemic body changes in the absence of a homeostatic mechanism (Gao, 2013). Thus, urine is a better source for biomarker samples. Urinary proteomics has received increasing attention in disease biomarker research.

Most of urinary proteins originate from the urinary system, and the remaining proteins are plasma proteins derived from glomerular filtration (Thongboonkerd and Malasit, 2005). Thus, the urinary proteome is expected to contain various biomarkers related to systemic disease. Currently, in addition to its extensive application in urogenital diseases, urinary proteomics have been applied to a wide range of non-urogenital diseases to exploit urinary disease biomarkers, such as cancers (Beretov et al., 2015), cardiovascular diseases (Brown et al., 2015), mental disorders (Wang et al., 2014), among others. However, challenges still remain in the urine biomarker discovery field. One important issue is that urine is a sensitive sample matrix. As summarized in a review (Wu and Gao, 2015), some physiological factors can influence the urine proteome, such as gender (Guo et al., 2015), age (Bakun et al., 2014), menstrual cycle (Castagna et al., 2011), exercise (Kohler et al., 2010), and smoking (Haniff and Gam, 2016). This sensitivity can impact the screening and subsequent validation of candidate disease biomarkers.

In clinical practice, drugs can reverse disease processes and are likely to have a significant impact on the urinary proteome and related disease biomarkers. However, due to ethical issues, it is impossible to halt drug treatment in patients during urine collection. Drugs have both therapeutic effects and side effects. Therapeutic effects cure the disease and might reduce or even eliminate disease biomarkers, whereas side effects create changes unrelated to the disease biomarker that might erroneously be considered disease biomarkers if the drug effect is not considered in a biomarker study. Knowing the impact of drugs on the urine proteome can help eliminate interference when detecting real disease biomarkers in urine.

As widely prescribed drugs, glucocorticoids are currently used to treat a wide range of diseases based on their potent anti-inflammatory, immunosuppressive and anti-neoplastic effects. One previous report indicated that approximately 0.9% of the population is using glucocorticoids at any given time (van Staa et al., 2000). Among glucocorticoids, prednisone is a commonly prescribed drug that can induce an overall catabolic protein response (Rose et al., 2010). Despite the high prevalence of prednisone use in clinical practice and its strong association with protein metabolism, the effects of prednisone treatment on the urinary proteome remain unknown, which might hamper biomarker discovery for numerous diseases. To evaluate the possible impact of prednisone administration on the urinary proteome and related disease biomarkers, this study performed urinary proteomics analysis using LC-MS/MS in a rat model. We found that prednisone treatment altered the urine proteome, and the pharmacological effects of prednisone were reflected in urine. Thus, urinary proteomics has the potential for drug efficacy monitoring in clinical practice. Meanwhile, changes to the urine proteome from prednisone treatment should be considered in future disease biomarker studies.

## Results and Discussion

### Characterization of prednisone-treated rats

To investigate the effect of prednisone treatment on the urine proteome, rats were treated with prednisone or saline for fourteen days. The body weight of the rats was measured each week. After analyzing the results, lower body weights were observed in rats receiving prednisone therapy compared with control rats on days 7 (245.83 ± 9.62 vs. 255.50 ± 7.04 g, prednisone vs. control) and 14 (279.50 ± 10.84 vs. 292.33 ± 9.27 g, prednisone vs. control). The six rats in the prednisone-treated group exhibited a slower increase in body weight than those in the control group throughout this experiment. However, this difference was not statistically significant (Figure 1A). Reduced body weight in the prednisone-treated rats might be due to growth inhibition caused by glucocorticoid treatment (Stikkelbroeck et al., 2003). After pathway analysis of differential proteins, it was found that the growth hormone signaling pathway significantly changed after prednisone treatment, which agrees with the reduced body weights observed in the prednisone group.

**Figure 1.**
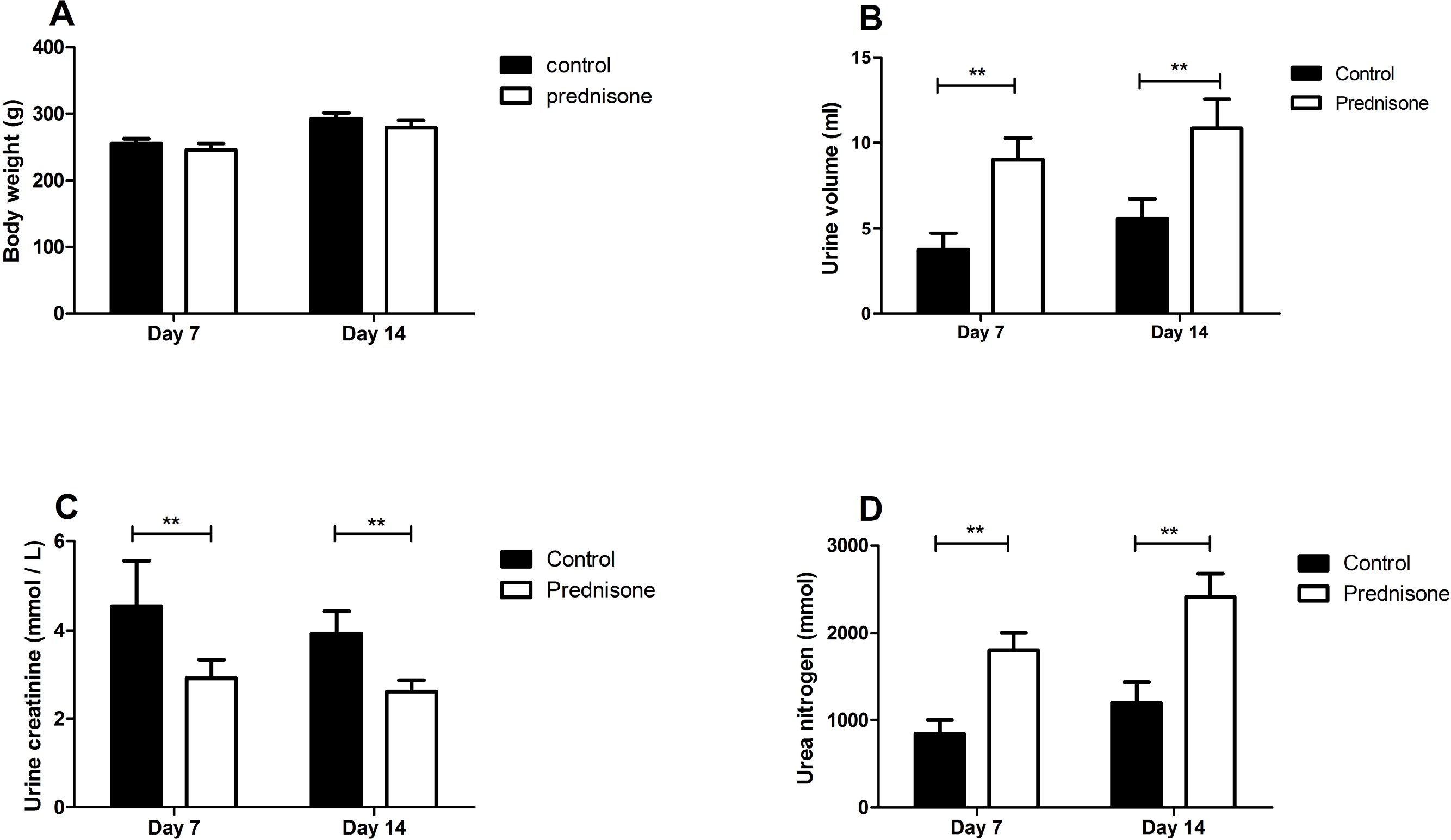
Body weights and urinary biochemical indicators in prednisone-treated and control rats. (A) The body weights of rats in the two groups on days 7 and 14. (B-D) Urine volumes, creatinine concentration and urea nitrogen levels in urine within 8 h of prednisone administration. The data represent the mean ± standard deviation (n = 6 in each group, **p < 0.01). Student’s t-test was used to assess whether the differences between the two groups were significant.

Meanwhile, we collected urine within 8 h of intragastric administration in the two groups, and urine volumes and biochemical indicators were measured. As shown in Figure 1B, rat urinary volumes increased significantly after prednisone treatment (3.75 ± 0.96 vs. 9.02 ± 1.25 ml, control vs. prednisone on day 7; 5.58 ± 1.67 vs. 10.85 ± 1.70 ml, control vs. prednisone on day 14, both p-values < 0.01). Because of the urine volume increase, the urine creatinine concentration was decreased in the prednisone group (Figure 1C, p < 0.05). The urine volume increase after prednisone treatment was consistent with the diuretic effect of glucocorticoids. In addition, as shown in Figure 1D, the excretion of urea nitrogen was significantly increased in urine from the prednisone group (843.21 ± 156.86 vs. 1800.69 ± 203.88 mmol, control vs. prednisone on day 7; 1,204.81 ± 233.58 vs. 2,417.41 ± 265.41 mmol, control vs. prednisone on day 14, both p-values < 0.01). Prednisone has been reported to induce an overall catabolic protein response and to accelerate protein breakdown (Rose and Herzig, 2013), leading the body to dispose of excess nitrogen waste as urea in urine. The strong association between prednisone treatment and protein metabolism might contribute to the changes in the urinary proteome. In summary, these results revealed that prednisone affected urine markers consistent with its pharmacological effects.

### Changes to the urine proteome after prednisone administration

For proteomics analysis, ten urine samples were randomly selected from the prednisone and control groups on day 14. Each sample was analyzed with two running times. Using the label-free quantification by the Progenesis LC–MS software, 523 urinary proteins were identified with a 1% false discovery rate (FDR) at the protein level. The criteria for identifying differentially expressed proteins included a fold change > 1.5 and p value < 0.05 between the two groups. Ultimately, 27 differentially expressed proteins were identified in urine with at least 2 peptides used for quantitation. Specifically, 12 proteins showed increased relative abundance, and 15 proteins showed decreased abundance (Table 1).

**Table 1:** Details of the identified urinary proteins altered by prednisone treatment.

| Uniprot | FC | Description | P | Human ortholog | IPA marker<sup>a</sup> | Urinary Protein Biomarker Database<sup>b</sup> |  |
| --- | --- | --- | --- | --- | --- | --- | --- |
|  |  |  |  |  |  | biomarker-animal | biomarker-human |
| P02764 | 1.77 | Alpha-1-acid glycoprotein | 0.004 | P02763 | Yes | ↑in DN | ↑in T1DM, DN, renal AR, AKI, acute appendicitis, vasculitis and pre-eclampsia |
| P05545 | 1.61 | Serine protease inhibitor A3K | 0.009 | P01011 | - | ↓in acute renal failure and nephrotic syndrome | ↑in T2DM, renal AR, non-small cell lung carcinoma and acute appendicitis |
| P09006 | 2.42 | Serine protease inhibitor A3N | 0.001 | P01011 | Yes | - | ↑in T2DM, renal AR, non-small cell lung carcinoma and acute appendicitis |
| P02780 | 2.66 | Secretoglobulin family 2A member 2 | 0.053 | - | - | ↓in nephrotic syndrome | - |
| P06866 | 2.08 | Haptoglobin | 0.047 | P00738 | Yes | ↑in nephrotic syndrome | ↑bladder transitional cell carcinoma, AKI, DN, and T2DM |
| P36374 | 2.01 | Prostatic glandular kallikrein-6 | 0.047 | - | - | ↓in acute renal failure | - |
| Q9EQV6 | 1.90 | Tripeptidyl-peptidase 1 | 0.001 | O14773 | - | - | - |
| O88766 | 1.69 | Neutrophil collagenase (MMP8) | 0.005 | P22894 | Yes | - | - |
| P02782 | 2.79 | Prostatic steroid-binding protein C1 | 0.038 | - | - | - | - |
| O54715 | 1.85 | V-type proton ATPase subunit S1 | 0.001 | Q15904 | - | - | - |
| Q9R0J8 | 1.51 | Legumain | 0.045 | Q99538 | - | - | - |
| P84039 | 2.11 | Ectonucleotide pyrophosphatase/phosphodiesterase family member 5 | 0.010 | Q9UJA9 | - | - | - |
| P23680 | -1.68 | Serum amyloid P-component | 0.006 | P02743 | Yes | ↑in anti-GBM nephritis and lupus nephritis | ↑in intestinal mucosal injury |
| P04764 | -1.57 | Alpha-enolase | 0.011 | P06733 | Yes | - | ↓in Dents disease |
| P81828 | -1.61 | Urinary protein 2 | 0.039 | - | - | ↓in nephrotic syndrome | - |
| P81827 | -1.68 | Urinary protein 1 | 0.038 | - | - | ↓in nephrotic syndrome and glomerulonephritis | - |
| P35444 | -1.62 | Cartilage oligomeric matrix protein | 0.011 | P49747 | Yes | ↑in osteoarthritic | - |
| P01830 | -1.90 | Thy-1 membrane glycoprotein | 0.036 | P04216 | Yes | - | ↑in prostate cancer |
| O35112 | -1.51 | CD166 antigen | 0.004 | Q13740 | Yes | - | ↑in T1DM |
| P52590 | -1.99 | Nuclear pore complex protein Nup107 | 0.007 | P57740 | - | - | ↓in IgA nephropathy |
| P15473 | -1.53 | Insulin-like growth factor-binding protein 3 | 0.017 | P17936 | Yes | - | ↑prostate cancer |
| P11030 | -1.56 | Acyl-CoA-binding protein | 0.029 | P07108 | - | - | ↑in Dents disease |
| P97603 | -1.50 | Neogenin (Fragment) | 0.009 | Q92859 | - | - | - |
| P80202 | -1.89 | Activin receptor type-1B | 0.046 | P36896 | - | - | - |
| P08649 | -1.66 | Complement C4 | 0.019 | P0C0L4 | - | - | - |
| P16310 | -1.58 | Growth hormone receptor | 0.039 | P10912 | - | - | - |
| Q5FVR0 | -1.51 | T-cell immunoglobulin and mucin domain-containing protein 2 | 0.020 | - | - | - | - |
| P14046 | -1.71 | Alpha-1-inhibitor 3 | 0.003 | P01023 | - | - | - |
| Q01205 | -1.61 | 2-oxoglutarate dehydrogenase complex component E2 | 0.026 | P36957 | - | - | - |

<sup>a</sup>Disease biomarkers filtered using IPA. <sup>b</sup>Urinary disease biomarkers from the Urinary Protein Biomarker Database identified in animal models and human diseases.
Abbreviations: T1DM: type 1 diabetes; T2DM: type 2 diabetes; DN: diabetic nephropathy; AKI: acute kidney injury; AR: allograft rejection

Several differentially expressed proteins identified in current study were confirmed to be regulated by glucocorticoids in previous studies, such as the serine protease inhibitor A3N (Nilsson et al., 2001), alpha-1-acid glycoprotein (Fournier et al., 1999), neutrophil collagenase (Muratore et al., 2009), growth hormone receptor (Vottero et al., 2003), insulin-like growth factor-binding protein 3 (Baxter, 1996) and acyl-CoA-binding protein (Compere et al., 2006), suggesting the reliability of the proteomics results. Meanwhile, we speculated that these urinary proteins were glucocorticoid responsive, similar to thrombospondin-1 (Barclay, et al., 2016), which has potential clinical utility as a glucocorticoid activity biomarker in urine.

Additionally, the measurement of 24 h urinary free cortisol (UFC) had been used to diagnose Cushing syndrome for more than four decades. However, the sensitivity and specificity of UFC are not ideal, particularly in the early stages of disease (Alexandraki and Grossman, 2011; Raff et al., 2015). Because exogenous glucocorticoid treatment can mimic hypercortisolism to some degree (Tavoni et al., 2013) and urine has the potential to reflect small and early changes in pathological conditions, our results suggested that specific urine proteins or protein patterns might be able to diagnose or screen for hypercortisolism (such as Cushing syndrome) as a supplement to urine cortisol testing. However, this should be verified in the urine of hypercortisolism patients.

### Functional analysis of urinary proteins

To identify major diseases and bio-functions of differentially expressed proteins involved, Ingenuity Pathway Analysis (IPA) Software was used for canonical function enrichment analysis. As shown in Figure 2, the principal diseases affected by the differentially expressed proteins included the inflammatory response, metabolic disease and cancer. The principal bio-functions of the identified proteins included cell death and survival, protein metabolism (degradation and synthesis), lipid metabolism, amino acid metabolism and carbohydrate metabolism. Prednisone is widely used in clinical practice based on its potent anti-inflammatory, immunosuppressive and anti-cancer effects. These results were consistent with the regulatory actions of prednisone in inflammation, immunity, metabolism and cancer, suggesting that the pharmacological effects of prednisone are reflected in the urine proteome after prednisone administration. As a non-invasive and sensitive biomarker source, urine has the potential to reveal many types of changes (pathological, pharmacological and physiological changes) in the body. In the future, the application of urinary proteomics will identify non-invasive biomarkers to detect drug toxicity or efficacy.

**Figure 2.**
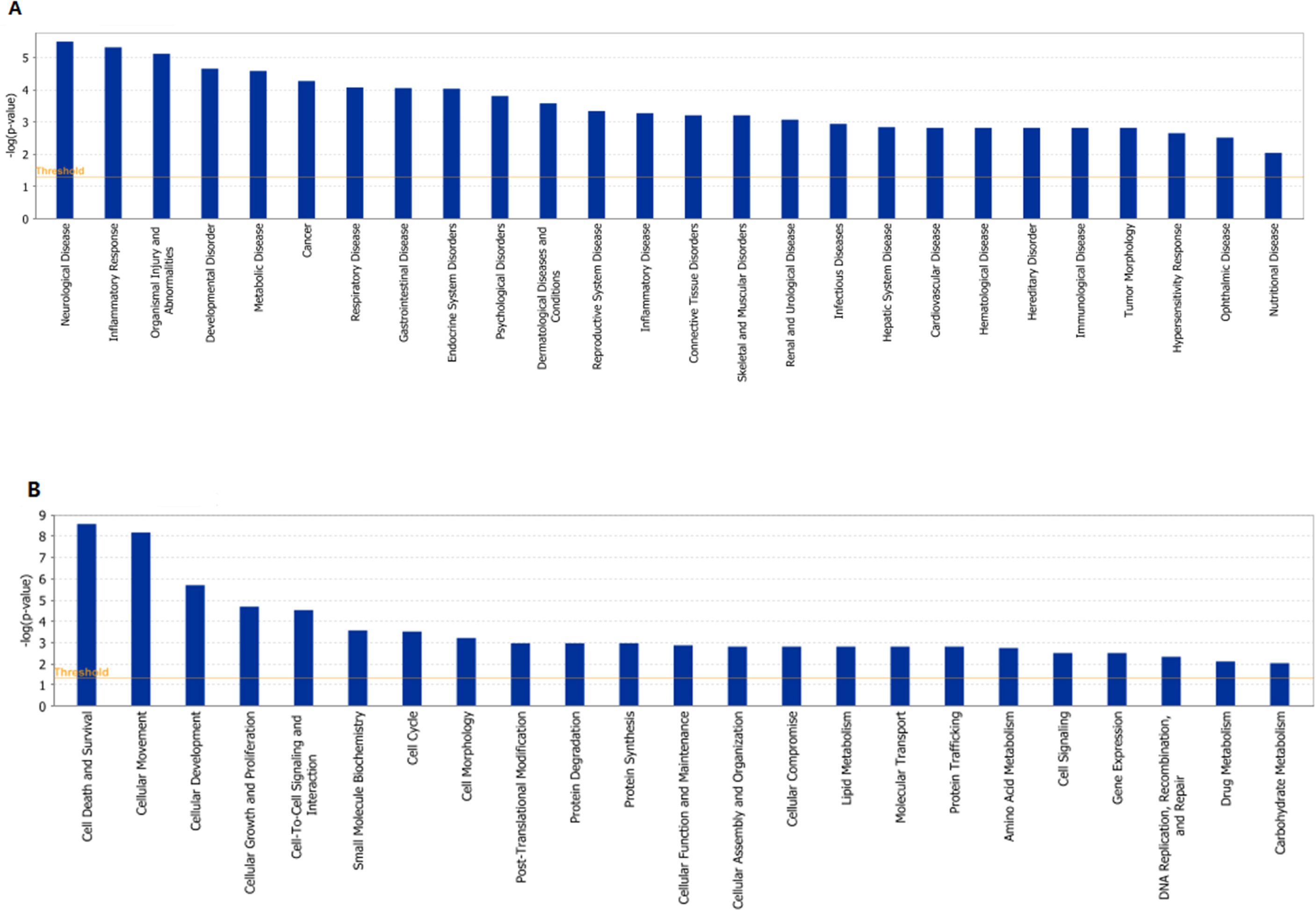
Analysis of principal diseases and bio-functions in which the differentially expressed proteins participate using IPA. Significance values were calculated based on Fisher’s right tailed exact test, and the –log(p-value) was calculated and is displayed on the y-axis of the bar chart. The taller the bar, the more significant the pathway effect.

### The effects of prednisone treatment on relevant candidate biomarkers in urine

Using the IPA biomarker filtering tool, 10 differentially expressed proteins were identified as candidate disease urine biomarkers. Because orthologs have similar bio-functions across different species, we converted the rat proteins to human proteins, and 23 rat urinary proteins had human orthologs. The Urinary Protein Biomarker Database is a literature curated database of protein biomarkers that have been detected in urine. When the selected differentially expressed proteins and their human orthologs were searched against the Urinary Protein Biomarkers Database (http://www.urimarker.com/biomarker/web/indexdb), 16 proteins were annotated as candidate urine biomarkers in previous studies. Integrating the results identified from these two databases, 16 proteins were considered to be potential disease biomarkers in urine. The expression trends of these biomarkers in various disease conditions are shown in Table 1.

Notably, the expression of some candidate biomarkers were completely reversed after prednisone treatment compared with their expression in specific pathological conditions. These proteins might serve as candidate therapeutic biomarkers for monitoring treatment effectiveness or reflecting glucocorticoid sensitivity. For example, up-regulated serum amyloid P (SAP) in the urine was identified as a biomarker of anti-glomerular basement membrane disease and lupus nephritis (Wu et al., 2010), and increased cartilage oligomeric matrix protein (COMP) in urine was observed in osteoarthritis (Misumi et al., 2006). Both proteins were downregulated in urine after prednisone treatment in our study; thus, these proteins might indicate the effectiveness of glucocorticoids in treating these diseases. Thus, previous studies of patients receiving drug treatments might have underestimated or not identified changes to candidate disease biomarkers. By contrast, the expression of some candidate biomarkers in our study after prednisone treatment were similar to levels in specific disease processes. These changes might not be relevant to the treatment effects of prednisone for these diseases and might even be erroneously considered disease biomarkers if the drug effect was not considered.

One important issue in the use of urine to detect biomarkers is that many factors can affect the urine proteome. Previous studies demonstrated that several common physiological factors influence the urine proteome. Compared with physiological factors, pharmacological agents are more important factors that should also be taken into consideration. In clinical practice, drugs can reverse the disease process and are very likely to have a significant impact on the urinary proteome and related disease biomarkers. Of note, physiological factors can often be well-matched for biomarker research. However, due to ethical issues, it is impossible to halt drug treatment in patients during urine collection or to give healthy volunteers drugs they do not need. Knowing the impact of drugs on the urine proteome can help us eliminate interference when detecting real disease biomarkers in urine. In previous studies, anticoagulants (Li et al., 2014) and the α1-adrenergic receptor antagonist (Zhao et al., 2016) were reported to influence the urine proteome and related biomarkers. Likewise, in the current study, 36 proteins were differentially expressed after prednisone treatment, and 17 of these proteins were previously reported candidate disease biomarkers. Two differentially expressed proteins (Haptoglobin [HPT] and Neutrophil collagenase [MMP8]) identified in our proteomic study with commercially available antibodies were selected to be validated by semi-quantitative western blot analysis (Figure 3). The changes in both proteins were consistent with the LC-MS/MS results.

**Figure 3.**
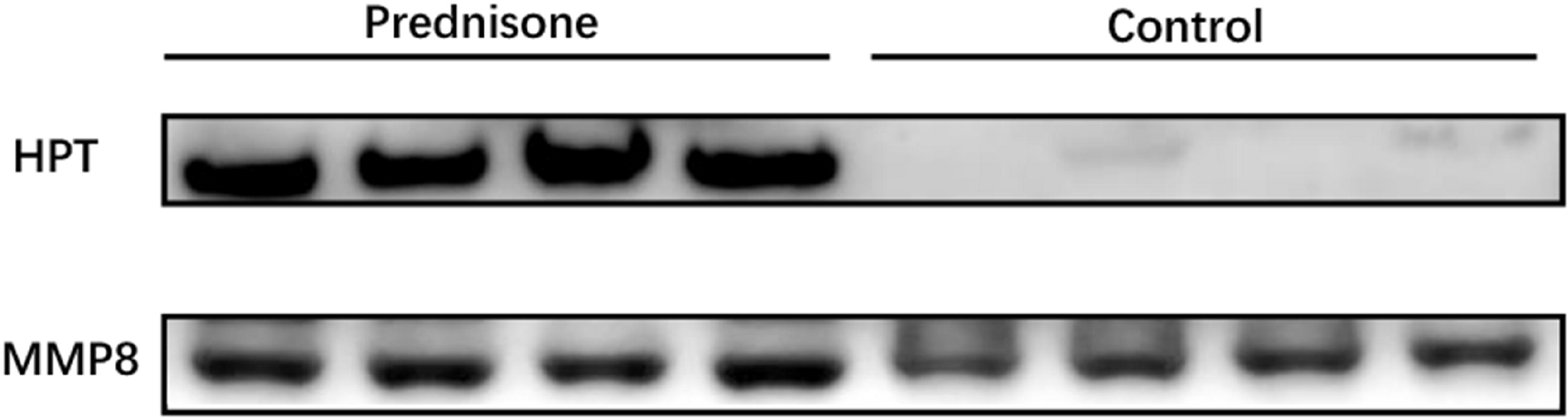
Western blot analysis of HPT and MMP8 in the urine samples after prednisone administration. A total of eight samples were used for validation (n = 4 in each group).

Prednisone is a widely prescribed drug in clinical practice, and its effects are strongly associated with protein metabolism. The significant changes to protein abundance identified by mass spectrometry suggested a large impact of prednisone on urine proteome and related urinary biomarkers. In future biomarker research, the changed protein pattern of urinary proteins in this study makes it possible to consider interference from prednisone. In addition, the functional analysis of the differentially expressed proteins suggests that the urinary proteome might reflect the pharmacological effects of prednisone. Urine is a sensitive and noninvasive sample resource, and the application of urine proteomics to drug monitoring may accelerate drug development and biomarker screening for drug toxicity or efficacy.

## Materials and Methods

### Experimental animals and drug administration

This study was performed on twelve male Sprague-Dawley rats (approximately 200 g) obtained from the Institute of Laboratory Animal Science, Chinese Academy of Medical Science. The animals were housed in cages with six rats per cage, fed a standard laboratory diet, and provided water ad libitum. All of the rats were housed under controlled indoor temperature (22 ± 1°C) and humidity (65 – 70%) conditions. All of the experimental protocols were reviewed and approved by Institute of Basic Medical Sciences Animal Ethics Committee, Peking Union Medical College (ID: ACUC-A02-2014-007).

After acclimatization for 3 days in cages, the rats were randomly divided into two groups. In the prednisone-treated group, rats were daily administered a prednisone solution (2 mg/ml) at 4 mg/kg/day via oral gavage. In the control group, rats were given a matching volume of sterile saline by intragastric administration. The rats in the prednisone group received medication every 24 h for 2 weeks. The animals were weighed every week, and the prednisone dose was adjusted based on changes in body weight.

### Urine collection and sample preparation

Urine samples were collected from all of the rats on days 7 and 14 after the animals were individually placed in metabolic cages. The rats had free access to water but no food to avoid sample contamination during urine collection. Urine was collected within 8 h of intragastric administration, and the urine volumes were measured. After collection, the urine was immediately centrifuged at 12,000 g for 20 min at 4°C. Urinary proteins were extracted with ethanol overnight followed by centrifugation at 12,000 g for 20 min. The precipitate was then resuspended in lysis buffer (8 M urea, 2 M thiourea, 50 mM Tris and 25 mM DTT). Sample aliquots were stored at −80°C for later proteomics analysis.

The urine after gavage was used for subsequent laboratory biochemical analysis. The creatinine and urea nitrogen concentrations in the urine were measured at the clinical laboratory of the Peking Union Medical College Hospital.

### Protein digestion

Urine samples from ten rats in two groups after gavage on day 14 were randomly selected (5 rats in each group). The urinary proteins were prepared using the filter-aided sample preparation method (Wisniewski et al., 2009). Briefly, after 100 µg of protein from an individual sample was denatured with 20 mM dithiothreitol at 37°C for 1 h and alkylated with 50 mM iodoacetamide in the dark for 40 min, the samples were loaded onto filter devices with a cut-off of 10 kD (Pall, Port Washington, NY, USA) and centrifuged at 14,000 g at 18°C. Then, after washing twice with UA (8 M urea in 0.1 M Tris-HCl, pH 8.5) and four times with 25 mM NH4HCO3, the samples were re-dissolved in 25 mM NH4HCO3 and digested with trypsin (enzyme to protein ratio of 1:50) at 37°C overnight. The peptide mixtures were desalted and dried by vacuum evaporation.

### LC-MS/MS analysis

The peptide samples were re-dissolved in 0.1% formic acid to a concentration of 0.5 µg/µL. For analysis, the peptides were loaded on a trap column and were separated on a reverse-phase analytical column using the EASY-nLC 1200 HPLC system. Then, the peptides were analyzed with an Orbitrap Fusion Lumos mass spectrometer (Thermo Fisher Scientific, Bremen, Germany). The elution for the analytical column was over 60 min at a flow rate of 300 nL/min. The mass spectrometer was set in positive ion mode and operated in data-dependent acquisition mode with full MS scan from 150 to 2,000 m/z with resolution at 120,000 and MS/MS scan from 110 to 2,000 m/z with resolution at 30,000. Two technical replicate analyses were performed for each individual sample.

### Label-free quantification

Label-free quantitation of the proteomic data was performed using Progenesis LC-MS software (version 4.1, Nonlinear, Newcastle upon Tyne, UK). Twenty sample features (ten samples with two technical replicates) were aligned according to their retention times, and peptides with charge states of +2 to +4 were selected in the analysis. The peak lists were exported, and the data were searched against the Swiss-Prot rat database (Released in July 2014; containing 7,906 sequences) using Mascot software (version 2.5.1, Matrix Science, London, UK). The parent ion tolerance was set at 10 ppm, and the fragment ion mass tolerance was set to 0.05 Da. A maximum of two missed cleavage sites in the trypsin digestion was allowed. Carbamidomethylation of cysteines was set as a fixed modification, and the oxidation of methionine was considered a variable modification. For protein quantification, the total cumulative abundance of a specific protein was calculated by summing the individual abundances of unique peptides. Comparisons across different samples were performed after normalization of protein abundance using Progenesis LC-MS software.

### Bioinformatics analysis and biomarker filtering

After label-free quantitation, the differential proteins were screened for a fold change > 1.5 and p value < 0.05 between the two groups. The differentially expressed proteins were further analyzed using IPA software (Ingenuity Systems, Mountain View, CA). This analysis was used to interpret the differentially expressed proteins based on the canonical pathways, interaction networks and disease mechanisms that the proteins were expected to regulate.

The biomarker filter function in the IPA software was used to filter disease biomarkers. Additionally, we identified the human orthologs of the differentially expressed proteins using BLAST (http://www.uniprot.org/blast/). These differentially expressed proteins and their human orthologs were searched using the Urinary Protein Biomarker Database (http://www.urimarker.com/biomarker/) to identify whether they were previously identified as candidate urinary disease biomarkers (Shao et al., 2011).

### Western Blot

Thirty micrograms of urinary protein from each sample (n=4 in each group) was loaded onto 10% SDS-PAGE gels and transferred to PVDF membranes with a transfer apparatus (Thermo Scientific Pierce G2 Fast Blotter). After blocking in 5% milk for 1 h, the membranes were incubated with primary antibodies overnight at 4°C. The primary antibodies used for validation included Haptoglobin (HPT) and Neutrophil collagenase (MMP8) (Abcam, USA). The membranes were then washed in TBST four times and incubated with secondary antibodies diluted 1:5,000 in a 5% milk solution for 1 h at room temperature. The immunoreactive proteins were visualized using enhanced chemiluminescence reagents (Thermo Scientific, USA). The protein signals were scanned with an ImageQuant 400TM Imager (GE Healthcare Life Sciences, Piscataway, NJ, USA), and quantified using the AlphaEaseFC system.

### Statistical analysis

Statistical analysis was performed with the Statistical Package for Social Studies (PASW statistics, SPSS, version 18.0). Data for the body weight, urine volume, and biochemical indicators in urine were presented as the mean ± standard deviation. All of the parameters were tested for normalization, and comparisons of these data between the control and prednisone group were performed using Student’s t-test. P-values of less than 0.05 were considered statistically significant.

## Compliance and ethics

The authors declare no conflict of interest. We conformed with the Helsinki Declaration of 1975 (as revised in 2008) concerning Human and Animal Rights, and followed out policy concerning Informed Consent as shown on Springer.com.

## Acknowledgments

This work was supported by the National Key Research and Development Program of China (grant number 2016YFC1306300), the National Basic Research Program of China (grant number 2013CB530850), and funds from Beijing Normal University (grant number 11100704, 10300-310421102).

